# Translational activation by a synthetic PPR protein elucidates control of *psbA* translation in Arabidopsis chloroplasts

**DOI:** 10.1101/2024.02.05.578914

**Authors:** Margarita Rojas, Prakitchai Chotewutmontri, Alice Barkan

## Abstract

Translation initiation on *psbA* mRNA in plant chloroplasts scales with light intensity, providing its gene product, D1, to replace photodamaged D1 in Photosystem II. The *psbA* translational activator HCF173 has been hypothesized to mediate this regulation. HCF173 belongs to the short-chain dehydrogenase/reductase superfamily, associates with the *psbA* 5’-untranslated region (5’-UTR), and has been hypothesized to enhance translation by binding an RNA segment that would otherwise pair with and mask the ribosome binding region. To test these hypotheses, we examined whether a synthetic pentatricopeptide repeat (sPPR) protein can substitute for HCF173 when bound to the HCF173 binding site. We show that an sPPR designed to bind HCF173’s footprint in the *psbA* 5’-UTR bound the intended site *in vivo* and partially substituted for HCF173 to activate *psbA* translation. However, sPPR-activated translation did not respond to light. These results imply that HCF173 activates translation, at least in part, by sequestering the RNA it binds to maintain an accessible ribosome binding region, and that HCF173 is also required to regulate *psbA* translation in response to light. Translational activation can be added to the functions that can be programmed with sPPR proteins for synthetic biology applications in chloroplasts.

**One sentence summary:** A synthetic PPR protein substitutes for HCF173, a non-PPR translational activator in chloroplasts, elucidating HCF173 functions and demonstrating the ability of synthetic PPRs to activate translation.

## Introduction

Control of gene expression at the level of translation offers the opportunity for rapid adaptation to environmental perturbations. This feature is particularly relevant for genes with long-lived mRNAs. Accordingly, translational control is much more prominent in chloroplasts than in their cyanobacterial ancestor, correlating with the much longer half-life of chloroplast mRNAs (Germain et al., 2013; Szabo et al., 2020; Wanney et al., 2023). Despite this difference, the basic mechanisms of translation in chloroplasts retain strong resemblance to those in bacteria. For example, chloroplast ribosomes initiate at internal ribosome binding sites characterized by unstructured RNA proximal to start codons, often with the aid of a Shine-Dalgarno element (Scharff et al., 2011; Scharff et al., 2017; Zoschke and Bock, 2018). However, unlike bacterial mRNAs, most chloroplast mRNAs require gene-specific translational activators to be translated efficiently. These translational activators are encoded by nuclear genes that evolved after the acquisition of the cyanobacterial endosymbiont. The vast majority are helical repeat proteins from the pentatricopeptide repeat (PPR), octotricopeptide repeat (OPR), or half-a-tetratricopeptide repeat (HAT) families (Rahire et al., 2012; Barkan and Small, 2014; Hammani et al., 2014; Ozawa et al., 2020). Current data support the view that these helical repeat activators stimulate translation by sequestering the RNA they bind, thereby preventing the formation of RNA structures that mask the ribosome binding site (Klinkert et al., 2006; Prikryl et al., 2011; Hammani et al., 2012; Rahim et al., 2016; Higashi et al., 2021). This protein-based mechanism mimics the RNA-based translational regulation by riboswitches and sRNAs in bacteria.

The chloroplast *psbA* mRNA offers a striking example of translational control in response to an environmental perturbation. The *psbA* gene encodes the D1 reaction center protein of Photosystem II. D1 is subject to photooxidative damage and the degree of damage scales with light intensity. Damaged D1 is removed by proteolysis, and must be replaced with newly synthesized D1 to maintain photosynthesis (reviewed in Jarvi et al., 2015; Theis and Schroda, 2016). The coupling of D1 synthesis to D1 damage in plant chloroplasts involves regulation at the level of translation initiation: the average number of ribosomes bound to each *psbA* mRNA molecule decreases dramatically within 30 minutes of shifting maize or Arabidopsis seedlings from moderate light intensities to the dark, and is restored after 15 minutes of reillumination (Chotewutmontri and Barkan, 2018). More subtle changes in *psbA* ribosome occupancy accompany a shift from low intensity to high intensity light in tobacco (Schuster et al., 2019). The light-induced signal that triggers these effects is D1 damage itself (Chotewutmontri and Barkan, 2020).

A thylakoid membrane complex involved in the initial steps of D1’s assembly into PSII is central to the autoregulatory mechanism that couples *psbA* translation to light-induced D1 damage (Chotewutmontri and Barkan, 2020; Chotewutmontri et al., 2020). This complex presumably relays a signal to factors bound to *psbA* mRNA that directly impact translation initiation. A protein called HCF173 is a strong candidate for serving in this role: in a comprehensive search for proteins that associate with *psbA* mRNA *in vivo*, HCF173 emerged as the only protein that binds *psbA* mRNA and is also required for its translation (Schult et al., 2007; McDermott et al., 2019; Watkins et al., 2020). Unlike most chloroplast translational activators, HCF173 is not a helical repeat protein. Instead, it consists of two globular domains: a short-chain dehydrogenase/reductase domain (SDR) and a CIA30 domain (see UniProt Q8W4D6). RNA coimmunoprecipitation data showed that HCF173 associates *in vivo* with an RNA segment in the *psbA* 5’-untranslated region (UTR) that maps adjacent to the sequence bound by the initiating ribosome (McDermott et al., 2019). Binding of that RNA by HCF173 was predicted to prevent formation of an inhibitory RNA structure involving the translation initiation region (see Figure 1A), suggesting a plausible mechanism by which HCF173 could activate translation (McDermott et al., 2019). This hypothesis is concordant with evidence that an increase in light intensity is accompanied by a decrease in RNA structure near the *psbA* start codon (Gawronski et al., 2021).

**Figure 1.**
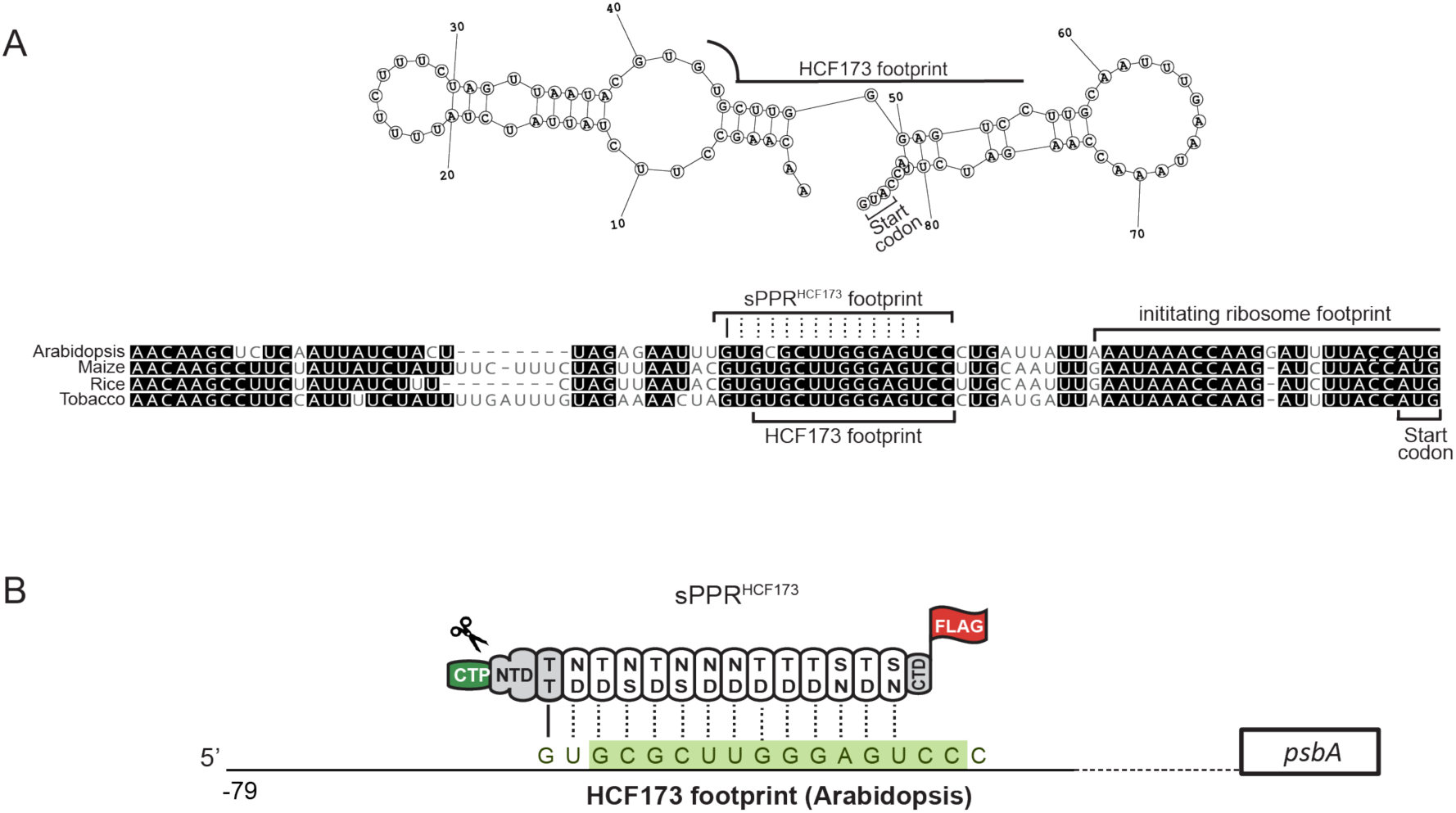
Experimental Design. (A) A multiple sequence alignment of the *psbA* 5’-UTR is annotated to indicate the footprint of HCF173, the footprint of the initiating ribosome, and the sequence to which sPPR^HCF173^ was targeted. The predicted secondary structure shown above was published previously (McDermott et al., 2019) and is copied here with permission. The structure was predicted with Dynalign (Fu et al., 2014) using the maize and Arabidopsis sequences as input. The prediction for the maize sequence is shown but Dynalign predicts a similar structure for Arabidopsis, and the Arabidopsis structure is supported by experimental data (Gawronski et al., 2021). (B) Schematic of sPPR^HCF173^. sPPR^HCF173^ contains 13 consensus PPR motifs flanked by sequences from PPR10, following the design of (Shen et al., 2015). The amino acids that specify the nucleotide bound by each repeat (Barkan et al., 2012) are indicated, with amino acids at positions 5 and 35 (numbering convention of (Yin et al., 2013)) shown at top and bottom, respectively. The PPR10-derived regions include its N and C-terminal regions (NTD and CTD, respectively), chloroplast targeting sequence (CTP) and first PPR motif (grey), which has an atypical specificity code (solid line). A FLAG tag is appended to the C-terminus of the PPR10-derived sequence. The targeting sequence is cleaved after import into the chloroplast (scissors).

Here we address the mechanism by which HCF173 activates translation and whether it is involved in perception of light. To do so, we examined whether a designer RNA binding protein with an entirely different architecture can substitute for HCF173 when bound to the HCF173 binding site. For that purpose, we took advantage of the ability to engineer PPR proteins to bind a desired RNA sequence. PPR proteins consist of tandem degenerate repeats that adopt a helical hairpin fold, which stack to form an elongated surface that binds RNA (reviewed in Barkan and Small, 2014; Shen et al., 2016). PPR tracts bind RNA in a modular one repeat-one nucleotide fashion, typically with considerable sequence-specificity. The identities of the amino acids at two positions within each repeat play a major rule in specifying the nucleotide bound (Barkan et al., 2012; Takenaka et al., 2013; Yagi et al., 2013; Yan et al., 2019). That said, there is considerable degeneracy in the "PPR code", and natural PPR proteins typically have idiosyncratic features that program sequence specificities whose basis is not understood (e.g. Miranda et al., 2017; Rojas et al., 2018). However, synthetic PPR proteins (sPPRs) consisting of consensus PPR motifs that differ only in the specificity determining positions can be engineered to reliably bind specified RNA sequences (reviewed in McDowell et al., 2022; Kwok van der Giezen et al., 2023).

Previously, we targeted an sPPR to the 3’-UTR of *psbA* mRNA *in vivo,* and used it as an affinity tag to purify *psbA* ribonucleoprotein particles for proteome analysis (McDermott et al., 2019). Here, we used the analogous protein design, modifying the specificity determining amino acids to target the sequence in the *psbA* 5’-UTR that is occupied by HCF173 *in vivo.* We found that this protein, denoted sPPR^HCF173^, is capable of substituting for HCF173 to activate *psbA* translation. Thus, translational activation can be added to the suite of applications demonstrated for sPPR proteins in chloroplasts. Although the activation achieved by the sPPR was less robust than that afforded by HCF173, our data suggest that this was due to insufficient sPPR abundance to occupy more than a small fraction of the extremely large *psbA* mRNA pool. These results support the hypothesis that HCF173 activates translation simply by sequestering the RNA to which it is bound. Furthermore, the enhanced ribosome occupancy promoted by the sPPR did not respond to light, indicating that HCF173 is essential for mediating the effects of light on *psbA* translation.

## Results

### Design of sPPR^HCF173^

HCF173’s *in vivo* "footprint", the sequence that coimmunoprecipites with HCF173 in the presence of ribonuclease (McDermott et al., 2019), maps to a conserved sequence block a short distance upstream from the footprint of the initiating ribosome (Figure 1A). This corresponds with the footprint of an unidentified protein detected by its resistance to micrococcal nuclease (Gawronski et al., 2021). The predicted secondary structure shown above, which is supported by experimental data (Gawronski et al., 2021), underlies the hypothesis that HCF173 activates translation by preventing the RNA it binds from pairing with the ribosome binding region. Given that natural PPR proteins activate translation in this manner (Prikryl et al., 2011; Higashi et al., 2021), the hypothesis predicts that an sPPR protein bound to the HCF173 footprint would be capable of substituting for HCF173. This prediction requires that *psbA* is HCF173’s sole target of action, which we showed previously by ribosome profiling analysis of an *hcf173* mutant (Williams-Carrier et al., 2019).

To test this prediction, we designed sPPR^HCF173^ to occupy the sequence marked in Figure 1A. sPPR^HCF173^ is identical to the sPPR we used to bind the *psbA* 3’-UTR *in vivo* (McDermott et al., 2019) except that the specificity-determining amino acids in the PPR tract were modified to match the targeted sequence in the *psbA* 5’-UTR. sPPR^HCF173^ (Figure 1B, Supplemental Figure S1) consists of the N-terminal portion of maize PPR10 (Pfalz et al., 2009) (including its chloroplast targeting sequence) followed by 13 consensus PPR motifs, and capped by a short segment at PPR10’s C-terminus. The footprint of sPPR^HCF173^ on the RNA (i.e. the region blocked from interaction with other proteins or RNA) is expected to extend roughly three nucleotides beyond the nucleotides that are directly bound by its PPR motifs (Prikryl et al., 2011; Barkan et al., 2012). A 3XFLAG tag was added at the C-terminus to allow for protein detection. This construct was introduced into Arabidopsis plants that were heterozygous for a null *hcf173* allele and transformed progeny expressing sPPR^HCF173^ were identified by immunoblot analysis of hygromycin resistant plants.

### Expression of sPPR^HCF173^ Restores Autotrophic Growth of *hcf173* Mutants

Due to their severe loss of PSII, *hcf173* mutants die shortly after germination unless supplied with exogenous sugar (Schult et al., 2007). By contrast, *hcf173* mutants expressing sPPR^HCF173^ (*hcf173:* sPPR^HCF173^) survived in soil through flowering (Figure 2A), and produced viable seeds. However, these plants grew more slowly than the wild-type and exhibited elevated chlorophyll fluorescence indicating compromised photosynthesis. When grown on agar medium supplemented with sucrose, *hcf173:* sPPR^HCF173^ plants grew more rapidly and were darker green than the parental *hcf173* mutants; they were, however, less robust than the wild-type and exhibited high chlorophyll fluorescence (Figure 2B). These results show that sPPR^HCF173^ can substitute for HCF173 to some extent, but this "complementation" was incomplete.

**Figure 2.**
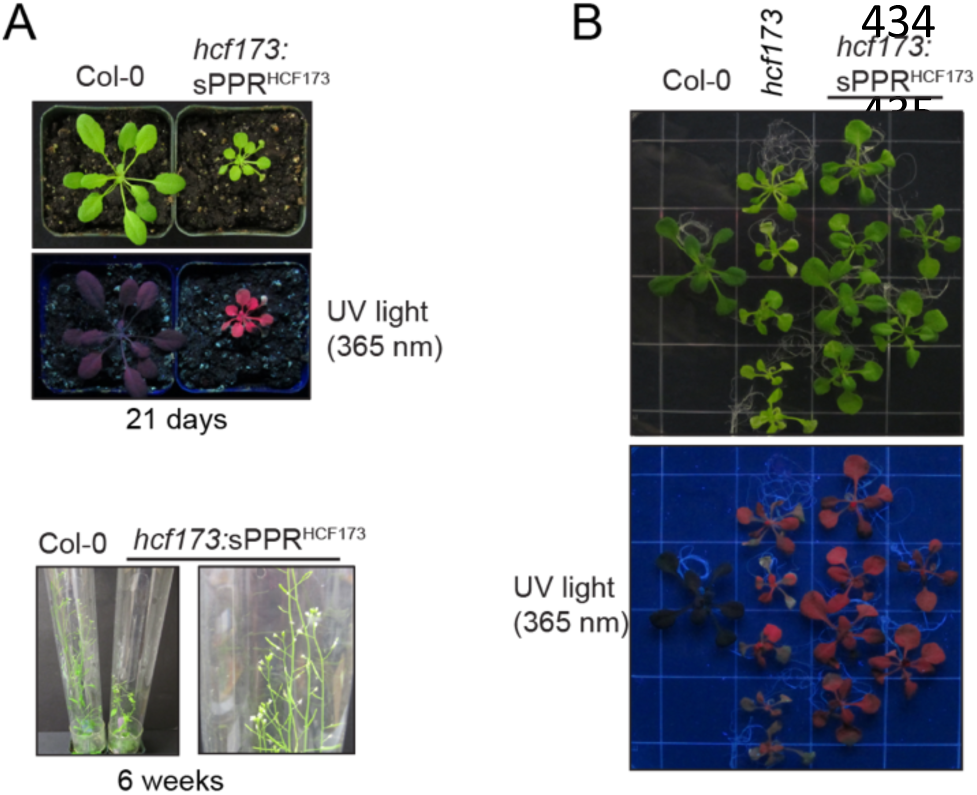
Phenotypes of *hcf173* mutants expressing sPPR^HCF173^. (A) Plants were sown on agar medium supplemented with 3% sucrose, transplanted to soil at two weeks and grown for an additional 1 week on soil. The same plants at flowering are shown below. *hcf173* mutants are not shown because they died 24 h after transplanting to soil. (B) Seedlings were grown for 19 days on agar medium supplemented with 3% sucrose.

### sPPR^HCF173^ Enhances the Translational Efficiency of *psbA* mRNA in *hcf173* Mutants

We next used immunoblot analysis to examine the effects of sPPR^HCF173^ on core subunits of photosynthetic complexes (Figure 3A). Whereas D1 was undetectable in *hcf173* mutants, it was readily detected in *hcf173:* sPPR^HCF173^ plants. However, D1 abundance in these plants remained well below that in the wild-type, consistent with their elevated chlorophyll fluorescence and slow growth. The abundance of the core PSI subunit PsaD was reduced in *hcf173* mutants; this is consistent with the PSI deficiency demonstrated previously in *hcf173* mutants, which was attributed to a secondary effect of the absence of PSII (Schult et al., 2007). Consistent with that view, amelioration of the PSII defect in *hcf173* mutants by sPPR^HCF173^ was accompanied by an increase in PsaD abundance (Figure 3A). Plastid-encoded subunits of Rubisco (RbcL), the cytochrome *b6f* complex (PetD), and ATP synthase (AtpB) were found at normal levels in *hcf173* mutants and were not affected by sPPR^HCF173^, suggesting that sPPR^HCF173^ did not have a deleterious effect on chloroplast gene expression in general.

**Figure 3.**
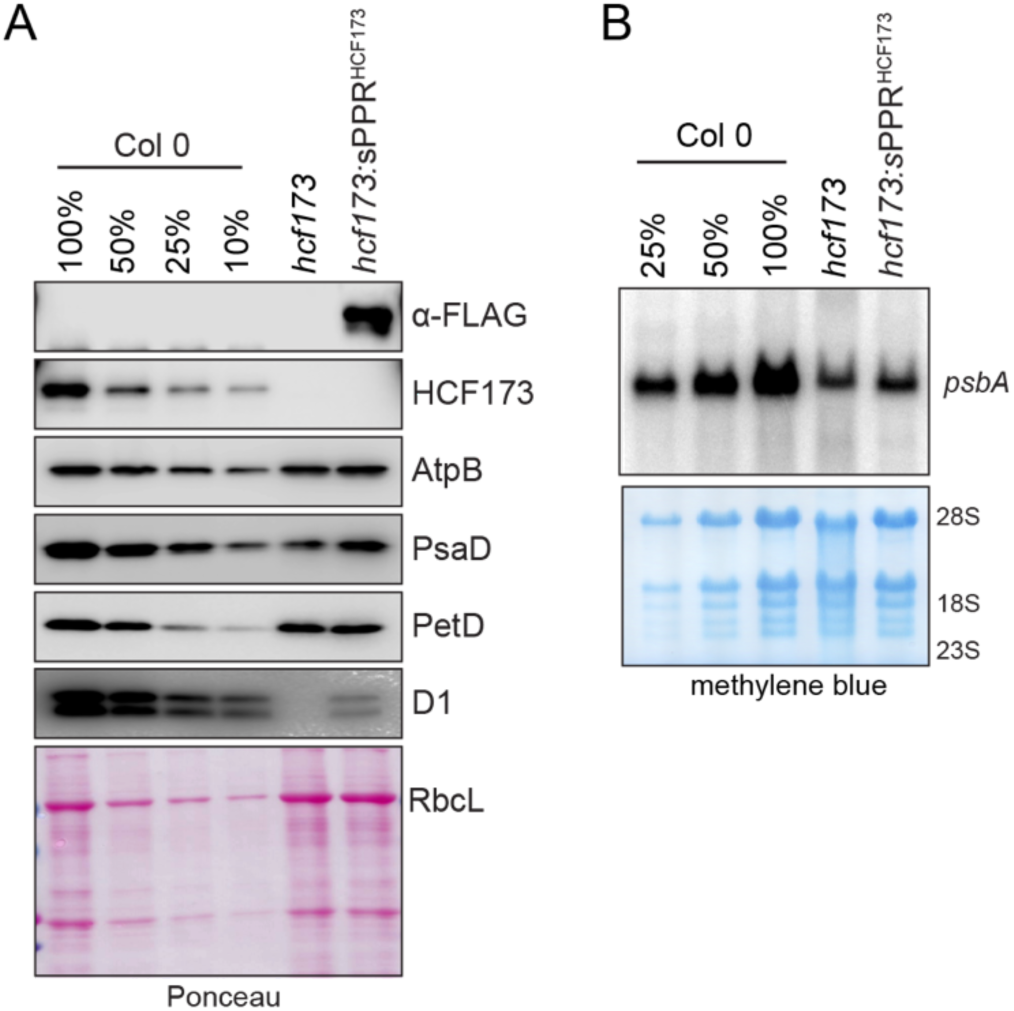
**Effects of sPPR^HCF173^ on the abundance of photosynthetic complexes and *psbA* mRNA.** (A) Immunoblot analysis of D1 and core subunits of ATP synthase (AtpB), cytochrome *b6f* (PetD), and PSI (PsaD). Probing for HCF173 confirmed the absence of HCF173 in the *hcf173* and *hcf173:sPPR^HCF173^* plants used for these experiments. sPPR^HCF173^ was detected with anti-FLAG antibody. Plants were grown for 19 days on agar medium supplemented with 3% sucrose. Replicate samples were run on two halves of the same gel, and the blot was divided for probing: one half was probed to detect HCF173, FLAG, and PsaD, and the other half was probed to detect D1, PetD and AtpB. The filter was stained with Ponceau S prior to probing to illustrate relative sample loading (below). (B) RNA gel blot hybridization of total leaf RNA illustrating reduced abundance of *psbA* mRNA in *hcf173* and *hcf173:*sPPR^HCF173^ plants. The blot was stained with methylene blue prior to probing (below), to illustrate relative sample loading.

The profound defect in *psbA* expression in *hcf173* mutants is due to the combined effects of decreased mRNA stability and decreased translational efficiency (Schult et al., 2007; Williams-Carrier et al., 2019). The presence of sPPR^HCF173^ caused little, if any, increase in the abundance of *psbA* mRNA in *hcf173* mutants (Figure 3B). We used ribosome profiling to examine the effect of sPPR^HCF173^ on the translational output of *psbA* and all other chloroplast genes. Ribosome footprints were prepared from leaves of seedlings that were dark-adapted for 1 hour or reilluminated for 15 minutes, and were analyzed by deep sequencing (Figure 4A, Supplemental Dataset S1). Previous RNA-seq analysis showed that the abundance of chloroplast mRNAs does not change detectably during the 15 minutes of reillumination (Chotewutmontri and Barkan, 2018), so changes in ribosome footprint abundance reflect changes in the average number of ribosomes per mRNA molecule. Consistent with prior results, the normalized abundance of ribosome footprints on *psbA* mRNA increased roughly six-fold during reillumination of Col-0, *psbA* translational output was reduced roughly 60-fold in reilluminated *hcf173* seedlings, and neither the light condition nor *hcf173* had a substantial impact on ribosome occupancy on other chloroplast mRNAs (Chotewutmontri and Barkan, 2018; Williams-Carrier et al., 2019). The presence of sPPR^HCF173^ increased the translational output of *psbA* roughly four-fold in *hcf173* mutants (compare *hcf173* to *hcf173:*sPPR^HCF173^ in Figure 4A), but this remained well below the output of *psbA* in the wild-type. Importantly, however, our data strongly suggest that the abundance of sPPR^HCF173^ was limiting for expression of *psbA*: the translational output of the nuclear transgene encoding sPPR^HCF173^ was quite variable among the individuals sampled for each replicate, and the translational output of *psbA* increased in proportion to the expression of sPPR^HCF173^ (Figure 4B, Supplemental Dataset S1). It is reasonable to extrapolate from these data that higher levels of sPPR^HCF173^ would restore *psbA* translation to a greater degree.

**Figure 4.**
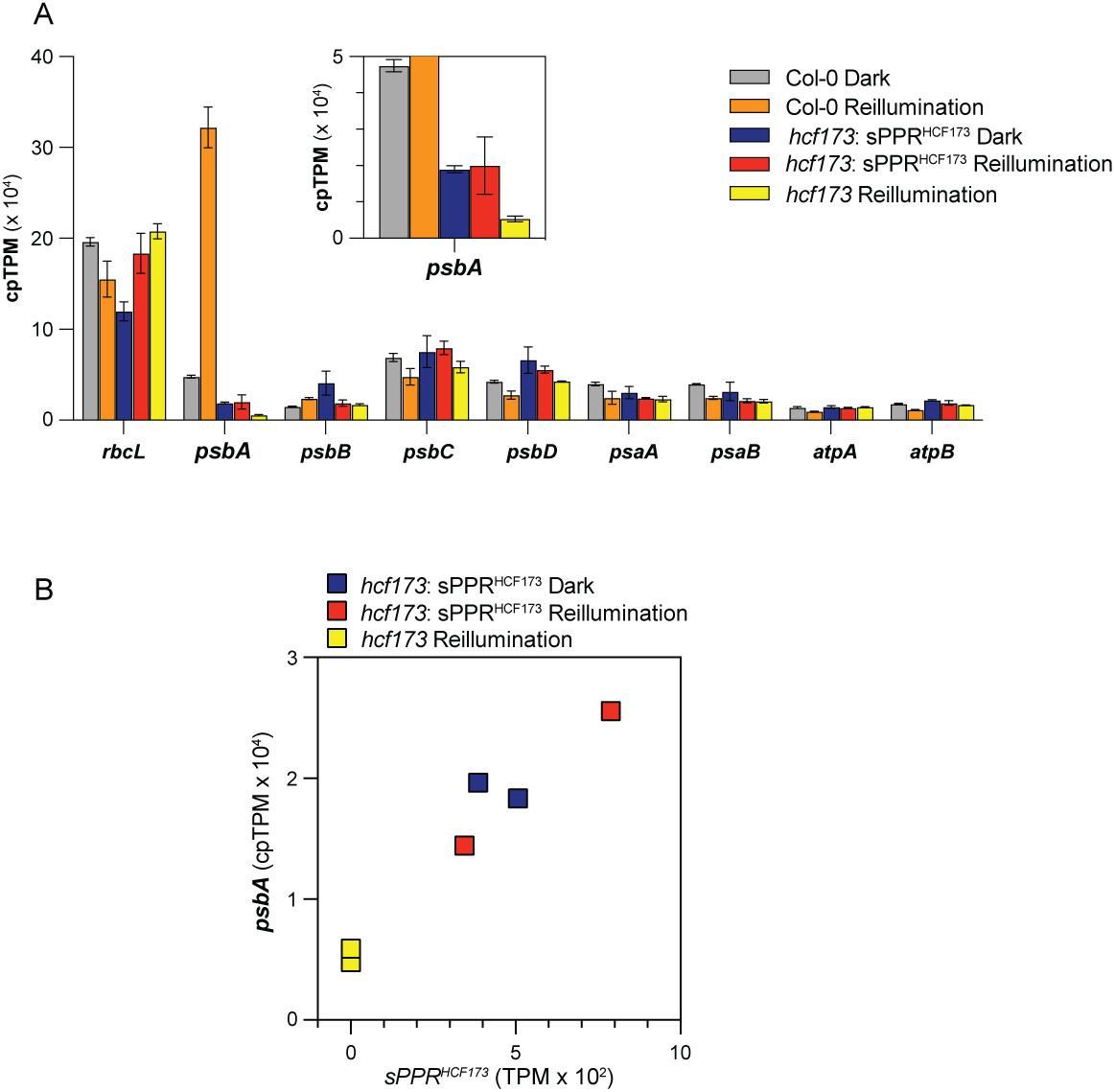
Ribosome profiling analysis of the effects of sPPR^HCF173^ on chloroplast gene output in dark-adapted and reilluminated plants. Plants were grown for 21 days on MS medium supplemented with 3% sucrose. Leaves of Col-0 and *hcf173:*sPPR^HCF173^ plants were harvested after 1 hour of dark adaptation at midday and after 15 minutes of reillumination. *hcf173* plants were harvested only after reillumination. Read counts were normalized to ORF length and sequencing depth using the TpM method (Transcripts Per Kilobase Million) (reviewed in Zhao et al., 2020). (A) Translational output of selected chloroplast genes (mean of two replicates +/- SD). Values for all genes are provided in Supplemental Dataset S1. (B) Correlation between translational output of *psbA* and translational output of the nuclear transgene encoding sPPR^HCF173^. Values are shown for each replicate individually.

### RNA Coimmunoprecipitation Confirmed the Association of sPPR^HCF173^ with the Targeted RNA Sequence and Revealed Off-Target Binding

The fact that sPPR^HCF173^ supported autotrophic growth of *hcf173* mutants and increased *psbA* translation strongly suggested that it bound the targeted sequence in the *psbA* 5’-UTR. To test this inference and to determine whether sPPR^HCF173^ exhibited off-target binding, we sequenced RNA that coimmunoprecipitates with sPPR^HCF173^ from the transgenic plants. In this "RIP-seq" experiment, whole leaf extracts from *hcf173* mutants expressing sPPR^HCF173^ were solubilized with non-ionic detergents, the cleared lysates were used for immunoprecipitations with anti-FLAG antibody, and the coprecipitating RNA was analyzed by deep sequencing. Parallel analysis of *hcf173* mutants lacking the transgene served as the negative control. Data from two replicate experiments (Dataset S2) are summarized in Figure 5A. The targeted sequence in the *psbA* 5’-UTR was represented by a well-defined plateau in read counts in the sPPR^HCF173^ data that corresponded precisely with the predicted footprint of sPPR^HCF173^ (Figure 5B, upper left). This sequence was enriched roughly 20-fold in immunoprecipitates from *hcf173:* sPPR^HCF173^ plants in comparison with *hcf173* mutants lacking the transgene (Dataset S2, Figure 5B). These results confirm that sPPR^HCF173^ bound *in vivo* to the intended sequence in the *psbA* 5’-UTR.

**Figure 5.**
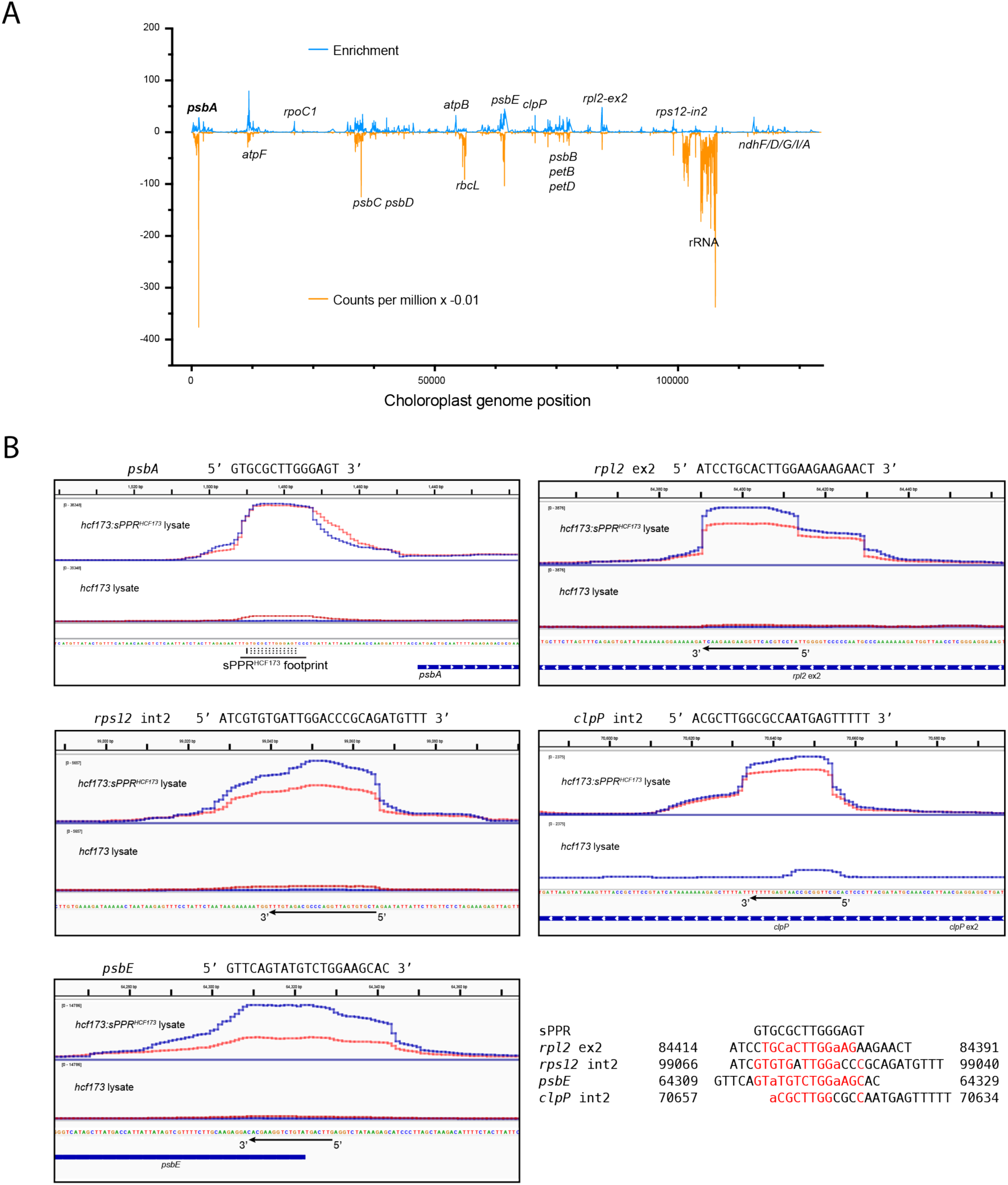
RIP-seq analysis of sPPR^HCF173^-RNA interactions *in vivo*. (A) Read counts (orange) and enrichment (blue) across the chloroplast genome. Only one copy of the large inverted repeat in the chloroplast genome is shown. Normalized read counts [counts per million reads mapped (CpM), average of two replicates of coimmunoprecipitation from *hcf173:*sPPR^HCF173^ lysate] are displayed for every position. Enrichment values (CpM *hcf173:*sPPR^HCF173^ / CpM *hcf173*) are not displayed for positions with fewer than two reads in each *hcf173* replicate to reduce spurious peaks resulting from low values in the denominator. Peaks corresponding to the intended target in *psbA* and to suspected off target sites are labeled above. Other major peaks are labeled below. Most of these encode integral membrane proteins, which were excluded from consideration as off-target sites due to the wide-spread enrichment of such mRNAs (see text). Values are provided in Supplemental Dataset S2. (B) Read coverage around the targeted sequence in *psbA* and at putative off-target sites. Candidate off-target sites were selected as regions with >500 read counts/million and >15-fold enrichment across at least 16 contiguous positions, excluding mRNAs encoding integral membrane proteins. Screen captures from the Integrative Genomics Viewer (IGV) show peak sequence coverage within those regions. Sequences within those peaks are aligned to the targeted RNA sequence at lower right. Similarities to the target RNA sequence are highlighted based on experimentally-determined degeneracies in the PPR code (Barkan et al., 2012; Miranda et al., 2018; Yan et al., 2019): U vs C mismatches are marked as matches (upper case red font) because these amino acid codes only weakly discriminate between the two pyrimidines; A residues at positions corresponding to G in the target sequence are marked as similarities (lower case red font), because G-binding motifs show substantial binding to A.

sPPR tracts bind well to numerous degenerate versions of their ideal RNA ligand, so some off-target binding is to be expected (Miranda et al., 2018; McDermott et al., 2019; Yan et al., 2019; Royan et al., 2021; Kwok van der Giezen et al., 2023). In fact, immunoprecipitation of sPPR^HCF173^ enriched RNA derived from many regions of the chloroplast genome (Figure 5A).

However, the widespread enrichment of mRNAs encoding proteins that are cotranslationally integrated into the thylakoid membrane (e.g. *atpF, psbC, psbD, petB)* suggests that incomplete solubilization of thylakoid membranes resulted in precipitation of membrane vesicles with bound polysomes translating *psbA*. Although some of these may represent direct off-target binding by sPPR^HCF173^, this complication precluded a comprehensive analysis of off-target sites. However, the strong enrichment of sequences derived from several mRNAs encoding soluble proteins (e.g. *rpl2)* or membrane proteins that are post-translationally integrated (*psbE*) (Zoschke and Barkan, 2015) is strongly suggestive of off-target binding by sPPR^HCF173^. The enrichment profiles of RNA sequences that emerged as top candidates are shown in Figure 5B. Each of these sites was represented by a plateau of sequence reads of similar breadth to that harboring the targeted sPPR^HCF173^, within which the sequence had considerable similarity to the site targeted by sPPR^HCF173^ (Figure 5B, lower right). These results strongly suggest that sPPR^HCF173^ bound to multiple off-target sequences *in vivo*. These interactions seem not to be of any physiological consequence, however, because the ribosome profiling data did not reveal a change in the expression of the corresponding genes.

## Discussion

The activation of *psbA* expression is essential for the repair process that maintains photosynthesis in the face of light-induced D1 damage (reviewed in Jarvi et al., 2015; Theis and Schroda, 2016). In angiosperm chloroplasts, this regulation occurs at the level of translation initiation (Chotewutmontri and Barkan, 2018, 2020). The nucleus-encoded protein HCF173 is a strong candidate for the factor that directly activates *psbA* translation in response to D1 damage because it is required specifically for *psbA* translation and it binds the *psbA* 5’-UTR (Schult et al., 2007; McDermott et al., 2019; Williams-Carrier et al., 2019). In this study, we advanced understanding of HCF173’s role in this phenomenon while also exploring the utility of synthetic PPR proteins as translational activators for synthetic biology applications.

We found that an sPPR protein that occupies the HCF173 binding site can substitute for HCF173 to activate *psbA* translation. Although the magnitude of the translational activation achieved in this experiment (maximum of five-fold) was much lower than that conferred by HCF173 itself, the fact that *psbA* translation scaled with expression of the sPPR^HCF173^ transgene showed that the concentration of sPPR^HCF173^ was limiting for *psbA* translation. This is not surprising given that *psbA* mRNA is the most abundant mRNA in leaf tissue, making up more than 25% of all mRNA molecules in the chloroplasts of Arabidopsis seedlings (Chotewutmontri and Barkan, 2018). Thus, the ability of sPPR^HCF173^ to activate translation when actually bound to the targeted RNA is likely to be considerably greater than we were able to demonstrate in this experiment.

The fact that sPPR^HCF173^ can substitute for HCF173 despite its entirely different architecture supports the simple translational activation mechanism proposed previously: sequestration of the RNA bound along the protein’s surface prevents that RNA from pairing with and masking the RNA that binds the initiating ribosome (McDermott et al., 2019). This general mechanism is well documented in chloroplasts, but all prior examples involve helical repeat proteins (Klinkert et al., 2006; Prikryl et al., 2011; Hammani et al., 2014; Rahim et al., 2016; Higashi et al., 2021), whose long RNA-binding surface makes them particularly well suited for this purpose. It remains possible, however, that HCF173 activates translation via multiple mechanisms, including, for example, interaction with the ribosome.

It had been unclear whether HCF173 is simply required for *psbA* translation or is also part of the regulatory mechanism that senses light-induced D1 damage. The severe *psbA* translation defect in *hcf173* mutants made this question difficult to answer by pulse-labeling, polysome analysis, or even ribosome profiling. By boosting *psbA* translation in *hcf173* mutants with sPPR^HCF173^, we were able to address this question. The fact that *psbA* ribosome occupancy did not change in response to light in *hcf173:*sPPR^HCF173^ plants strongly suggests that HCF173 is an essential component of the mechanism that senses light-induced D1 damage. HCF173’s unusual protein architecture, consisting of a discontinuous SDR domain interrupted by a CIA30 domain, seems likely to be integral to its signal transduction function.

Previous applications of synthetic PPR proteins include use as an affinity tag to immunopurify a specific chloroplast mRNP (McDermott et al., 2019), substitution for endogenous PPR RNA stabilization factors in chloroplasts (Manavski et al., 2021), regulation of splice site choice in mammalian cells (Yagi et al., 2022), substitution for an endogenous chloroplast PPR RNA editing factor (Royan et al., 2021) and editing of that same chloroplast RNA sequence when expressed in *E. coli* (Bernath-Levin et al., 2021). Our findings add translational activation to the repertoire of functions that can be programmed with sPPR proteins.

Translational activation is a well-documented function of natural PPR proteins in chloroplasts and mitochondria (Barkan et al., 1994; Yamazaki et al., 2004; Tavares-Carreon et al., 2008; Pfalz et al., 2009; Cai et al., 2011; Prikryl et al., 2011; Zoschke et al., 2012; Haili et al., 2016; Zoschke et al., 2016; Williams-Carrier et al., 2019), and modification of specificity determining amino acids in natural PPR translational activators can extend this function to a limited set of engineered 5’-UTRs (Rojas et al., 2019). The ability to use synthetic PPR proteins for this purpose expands the range of RNA sequences that can be used as translational regulatory elements for synthetic biology applications in chloroplasts.

Our data strongly suggest that sPPR^HCF173^ bound several RNAs in addition to its intended target. This was not unexpected, given that many native PPR proteins affect the metabolism of multiple RNAs harboring related *cis*-elements, as well as prior evidence for off-target interactions of synthetic PPR proteins in chloroplasts (McDermott et al., 2019; Royan et al., 2021) and *E. coli* (Bernath-Levin et al., 2021). Although the off-target interactions detected here had no apparent impact on chloroplast gene expression, the possibility for problematic off-target effects scales with the size of the transcriptome. This will be of particular concern for applications in the nucleo-cytosolic compartment. Our finding that sPPR^HCF173^ abundance was limiting for *psbA* translational activation was foreshadowed by a similar observation made with a synthetic "PLS"-type PPR RNA editing factor in chloroplasts (Royan et al., 2021), highlighting another challenge to be addressed. Thus, there is a need to optimize sPPR designs to increase protein stability and expression, while also decreasing the propensity for off target binding.

Recent advances in sPPR protein design, fostered by the development of high throughput screening methods (Bernath-Levin et al., 2021; Yagi et al., 2022), hold promise for overcoming these challenges.

## Methods

### Generation of Transgenic Plants Expressing sPPR^HCF173^

The transgene encoding sPPR^HCF173^ is identical to that of the transgene encoding SCD14 we used to immunopurify *psbA* mRNPs (McDermott et al., 2019) except that the specificity determining amino acids in the sPPR tract were modified to match the HCF173 binding site (see Fig. 1A). The construct employs the sPPR design of Shen et al. (Shen et al., 2015) which embeds an sPPR tract between the N- and C-terminal regions of the natural PPR protein PPR10 (Pfalz et al., 2009).

The sequence encoding sPPR^HCF173^ was codon optimized for Arabidopsis, synthesized by Genewiz (South Plainfield, New Jersey) and cloned into a modified version of pCambia1300 harboring a "Superpromoter" (Lee et al., 2007), such that a 3xFLAG tag was fused in-frame to the C-terminus of the protein (see (McDermott et al., 2019) for details). The vector was a generous gift of Jie Shen and Zhizhong Gong (China Agricultural University). Arabidopsis Col-0 plants were transformed by the floral dip method (Zhang et al., 2006) and progeny expressing the transgene were identified by resistance to hygromycin and immunoblot detection of sPPR^HCF173^ using anti-FLAG antibody. Plants expressing sPPR^HCF173^ were crossed with plants that were heterozygous for a null allele of *hcf173* (SALK_035984) (Williams-Carrier et al., 2019).

Progeny that were homozygous for *hcf173* were identified by PCR using primers 5’ agtaacatggctgcgactga and 5’ gtagccacggagcatgagtt flanking the T-DNA insertion, in conjunction with a primer reading outward from the T-DNA left border (5’ tggttcacgtagtgggccatcg 3’). The absence of HCF173 and presence of sPPR^HCF173^ were confirmed by immunobloung using antibodies to HCF173 and the FLAG tag, respectively.

### Plant Growth

Seeds were sterilized in 1% (v/v) bleach and 0.1% (w/v) SDS for 10 min, washed in 70% (v/v) ethanol, rinsed three times in sterile water, and plated on Murashige and Skoog (MS) agar medium (Sigma-Aldrich) supplemented with 3% (w/v) sucrose. Transgenic plants were selected by the addition of 200 µg/mL hygromycin to the growth medium. Seedlings were grown in a growth chamber in diurnal cycles (10 h of light at 100 µmol photons m^−2^ s^−1^, 14 h of dark, 22°C) for 21 d. Seedlings used for RNA gel blot hybridizations, immunobloung, ribo-seq and RIP-seq were harvested after roughly 3 weeks of growth; specifics for each experiment are provided below. Plants used for propagation were transferred to soil and grown in 16 h light (100 µmol photons m^−2^ s^−1^), 8 h dark cycles, 22°C).

### Immunoblot and RNA gel blot analyses

Protein was extracted from leaf tissue, resolved by SDS-PAGE and analyzed by immunobloung as described previously (Barkan, 1998) except that precast Tris-Glycine 4-20% polyacrylamide gels (Novex, Invitrogen) were used for gel electrophoresis. Antibodies to PsaD, AtpB, D1, and PetD were described in (McCormac and Barkan, 1999) and the antibody to HCF173 was described in (McDermott et al., 2019). Anti-FLAG (M2 clone) antibody was purchased from Sigma-Aldrich (catalog # F1804 1MG).

RNA was isolated from leaves of 19 day old seedlings using TRIzol Reagent (Invitrogen) and analyzed by RNA gel blot hybridization as described previously (Barkan, 1998) with modifications to the hybridization conditions due to the use of a synthetic oligonucleotide (IDT) as a probe for *psbA* mRNA (5’ ct tcg ci tcg cgt ctc tct aaa ai gca gtc at 3’). The probe was 5’-end labeled with [γ-^32^P]-ATP and T4 Polynucleotide Kinase (Thermo Fisher Scientific, Waltham, MA, USA) according to the manufacturer’s instructions. Prehybridization, hybridization, and washing were performed as described for (Zhelyazkova et al., 2012), and the blot was imaged on a Typhoon Imager (Amersham).

### Ribosome Profiling

Plants used for ribosome profiling were grown in diurnal cycles (10 h light at 100 µmol photons m^−2^ s^−1^, 14 h of dark, 22°C) for 21 days on MS medium supplemented with 3% sucrose. Plants were transferred to fresh media every week. Seed used for the *hcf173*:sPPR^HCF173^ planDng was pooled from the progeny of four plants that were homozyogous for *hcf173* and expressing similar levels of sPPR^HCF173^. The aerial porDon of five plants was pooled for each replicate analysis of *hcf173* and *hcf173*:sPPR^HCF173^. The aerial portion of two plants was pooled for each replicate of Col-0.

Ribosome footprint preparation and rRNA depletion were performed as described previously (Chotewutmontri et al., 2018). The NEXTFLEX Small RNA-Seq Kit v4 (Perkin Elmer) was used for library construction.

### RIP-seq

Plants used for RIP-seq (*hcf173* and *hcf173*:sPPR^HCF173^) were grown for 21 days under the same conditions as plants used for ribosome profiling, and were harvested by flash freezing in liquid nitrogen at midday. The aerial portion of 10 plants (∼300 mg fresh weight) was pooled for each replicate. Frozen tissue was ground in liquid nitrogen in a mortar and pestle in polysome extraction buffer (50 mM Tris-acetate pH 8.0, 0.2 M sucrose, 0.2 M KCl, 15 mM MgCl2, 2% polyoxyethylene (10) tridecyl ether, 1% Triton X-100, 20 mM 2-mercaptoethanol, 100 μg/mL chloramphenicol, 2 μg/ mL pepstatin, 2 μg/ mL leupeptin, 2 mM PMSF), allowed to thaw, and then cleared twice by centrifugation at 15,000 g at 4°C for 10 min. RNAsin (Promega) was added to the cleared lysate at 1.7 U/μL. Immunoprecipitation and RNA purification were performed as described previously (McDermott et al., 2019). In brief, anti-FLAG M2 antibodies (Sigma) were pre-bound to Pierce Protein A/G Magnetic Beads (Thermo Fisher Scientific) in co-immunoprecipitation buffer (20 mM Tris-HCl, pH 7.5, 150 mM NaCl, 1 mM EDTA, 0.5% (v/v) Nonidet P-40, and 5 µg/mL aprotinin), using sufficient antibody to capture all sPPR^HCF173^ protein in the lysate, as determined in pilot experiments. The beads were washed 4 times with the same buffer prior to addition to the cleared lysate. The mixture was incubated on a rotator at 4°C for 1.5 h, after which the beads were collected to the side of the tube with a magnet and washed 5 times with co-immunoprecipitation buffer. The washed beads were then resuspended in 0.6 mL of ribosome dissociation buffer (10 mM Tris-Cl, pH 8.0, 10 mM EDTA, 5 mM EGTA, 100 mM NaCl, 1% SDS) and RNA was isolated with Tri reagent (Molecular Research Center). Each sequencing library was prepared from 80 ng of immunoprecipitated RNA. The RNA was first fragmented by incubation in 40 mM Tris-acetate pH 8, 100 mM potassium acetate, 30 mM magnesium acetate at 95°C for 4 min. Fragmentation was stopped by addition of EDTA to 50 mM. The RNA was ethanol precipitated and phosphorylated with T4 polynucleotide kinase (New England Biolabs) prior to library construction using the NEXTFLEX Small RNA-Seq Kit v4 (Perkin Elmer). The libraries were gel purified to select for inserts between 15 and 100 nucleotides before sequencing.

### Illumina sequencing and data analysis

Sequencing was performed by the Genomics and Cell Characterization Core Facility at University of Oregon using a NovaSeq 6000 instrument in 118-nucleotide single read mode. Sequencing data were processed as described in (Chotewutmontri et al., 2020) except that all read lengths were included for RIP-seq analysis. Read counts mapping to chloroplast genes are provided in Supplemental Datasets S1 and S2 for ribo-seq and RIP-seq, respectively.

### Accession Numbers

The gene identification number for HCF173 is AT1G16720. The RIP-seq and ribo-seq data were submitted to the SRA database under BioProject number PRJNA1054207. These data will be released upon publication. A reviewer can access the read-only link here: https://dataview.ncbi.nlm.nih.gov/object/PRJNA1054207?reviewer=g87q0heltp7uk7s634ilrirlmm

## Acknowledgements

We are grateful to Rosalind Williams-Carrier for comments on the manuscript. This work benefited from access to the University of Oregon high performance computing cluster, Talapas. This work was funded by grants MCB-2034758 and IOS-2052555 to A.B. from the National Science Foundation.

---

**Supplementary Figure S1.**
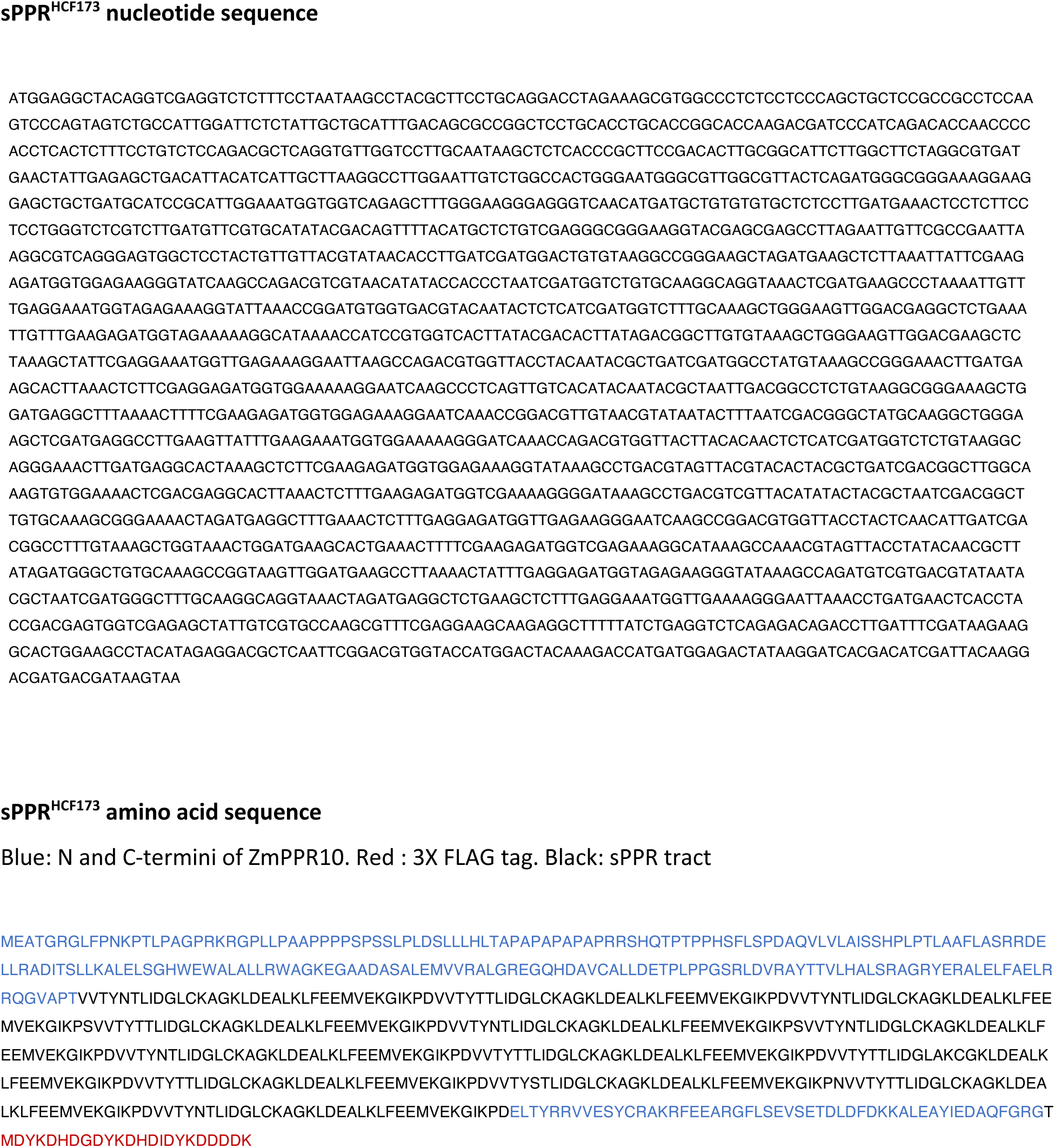
Sequence of sPPR^HCF173^ and the transgene encoding it.

**Table S1. Summary of ribo-seq data.**

**Table S2. Summary of RIP-seq data.**

